# Integration of mobile sequencers in an academic classroom

**DOI:** 10.1101/035303

**Authors:** Sophie Zaaijer, Columbia University Ubiquitous Genomics 2015 class, Yaniv Erlich

## Abstract

The rapid development of DNA sequencing technologies creates new educational opportunities for hands-on training. We report our experience in integrating handheld DNA sequencers (Oxford Nanopore Technologies’ MinION) as part of an academic class. This manuscript describes lessons learned to facilitate successful integration and provides educational resources for the benefit of the community.

## Main Text

The last decade has witnessed dramatic changes in the field of genomics with the advent of high-throughput sequencing technologies. Sequencers have become the ultimate apparatus for a wide array of applications, from genetic testing to forensic sciences. The data generated from these tasks form typical examples of “big data” science (Donovan 2008; Graham-rowe, Waldrop, and Lynch 2008). With such diverse applications, education in genome sciences can benefit from hands-on training in order to facilitate integrative thinking with respect to the technological, ethical, and scientific challenges (Altman 1998; Magana et al. 2014). Hands-on training is also the preferred learning style of the Millennial generation, which currently makes up the majority of undergraduate and graduate students. Research has shown that Millennials are technology focused, work most effectively in groups and absorb information most efficiently by kinesthetic learning (learning by doing) (Shapiro et al. 2013; Evans, Ozdalga, and Ahuja 2015; Linderman et al. 2015).

Here, we describe our experience with integrating mobile DNA sequencers in the classroom to facilitate hands-on learning. Our class focused on the newest sequencing technology: the MinION by Oxford Nanopore Technologies (ONT). Unlike other sequencing technologies that are static and require a laboratory setting, the MinION sequencer is slightly larger than a typical USB stick and only requires a laptop for sequencing (**Figure 1A** & **B**). It can be used at the office or in the field (Gardy, Loman, and Rambaut 2015; McIntyre et al. 2015; Erlich 2015). Some other distinct features include the following: 1) DNA libraries generated directly from genomic DNA; 2) sequence reads are long (up to 250 kb, in contrast to short Illumina reads); and 3) each DNA molecule that completes passage through the nanopore can be analyzed within minutes after a cloud-based base-calling step. The arrival of such handheld sequencers ushers in a new range of applications from at-home sequencing to new devices with DNA awareness (Erlich 2015). Here we describe our experience with teaching genomics by offering students mobile sequencers for hands-on learning. All relevant teaching material is provided under the Creative Commons Attribution-Share Alike 4.0 International License to facilitate use by the community.

**Figure 1:**
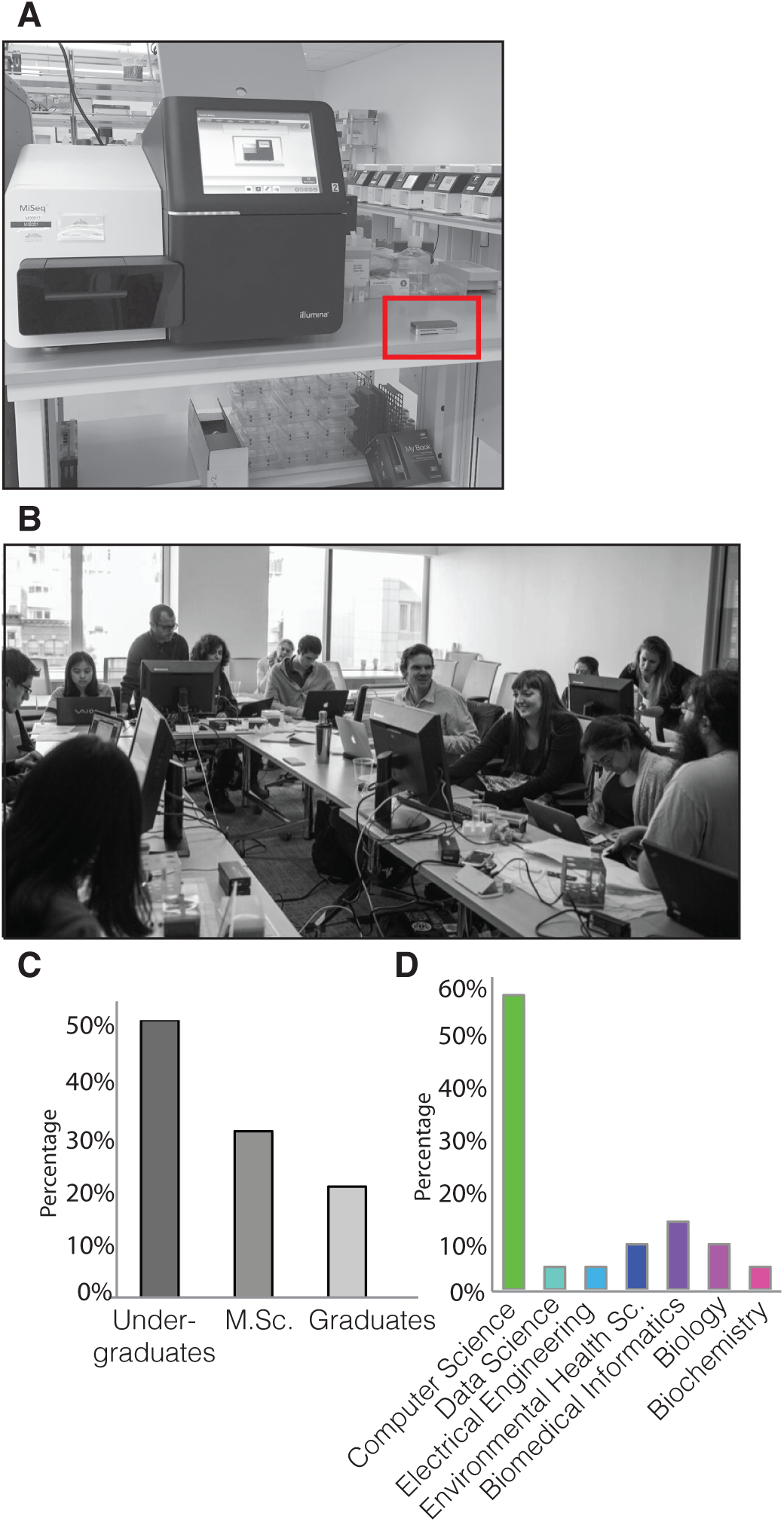
Overview of the Ubiquitous Sequencing class (A) Illumina MiSeq benchtop sequencer (left) versus MinION sequencer from ONT (right: red rectangle) (B) The hackathon class set-up (C)Distribution of student programs (D) Distribution of students degrees.

## Overview of the Ubiquitous Genomics class

We developed a course for Columbia University entitled ‘Ubiquitous Genomics’ that brings portable sequencing to the classroom. A total of 20 students enrolled in the course. Most of the students were studying towards Bachelors or Masters degrees (**Figure 1C**). The students’ degrees were variable, including computer science, electrical engineering, and biology (**Figure 1D**).

The course had 12 classes (one two-hour class per week) separated into two blocks. The first six-week block was an overview of sequencing technologies and various applications in medicine, bio-surveillance, and forensics. The aim of this block was to create a common ground for the group of students with such diversity in majors and background knowledge. The format was an active seminar where the class discussed one or two recent research papers. The second half of the course was devoted to hands-on learning and consisted of two three-week blocks of hackathons.

In the first week of each block, students met for a ~3 hour session of MinION sequencing. The second class in the block covered a specific technical area related to the sequencing experiment, such as alignment algorithms and involved active exploration and testing of the students. The final class of each hackathon block was devoted to student presentations of results (**Supplemental Note 1**).

We designed the hackathons to maximize the hands-on experience of the students within the time constraints of the class (**Figure 2**):

**Figure 2.**
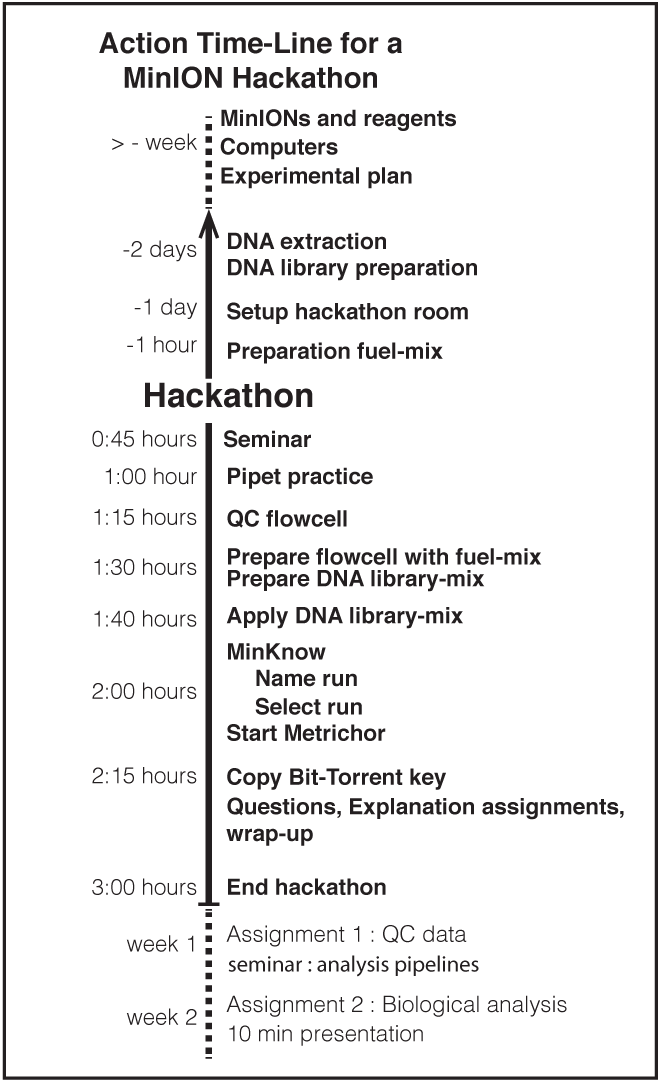
A detailed workflow for running a hackathon using MinION

A week before the hackathon the students were instructed to form groups of 4–5 people. We encouraged them to form groups with diverse backgrounds (e.g. a combination of biology majors and computer science majors).

We decided to prepare the DNA libraries several days before the hackathons and not to include them as part of the training (**Supplemental Note 2**). With the current technology, genomic DNA extraction and ONT library preparation takes ~5 hours and it was not realistic to include these steps as part of the hackathons (although this might change with the advent of the automated library preparation device; the VolTRAX).

Each hackathon started with a 45 min lecture about the goals of the hackathon, and some background material. We also gave an introduction to the ONT library preparation protocol, the software interface (MinKnow), and the base-calling pipeline (using Metrichor) in the lecture introducing the first hackathon (see **Supplemental Note 3–6** for assignments and PowerPoint slides).

Next, we had students practice pipetting. The loading of reagents onto the MinION flow cell requires good pipetting skills otherwise the yield may be substantially lower. As most of our students had never touched a pipet before, we allowed them to practice loading water onto used MinION flow cells until they were comfortable pipetting with precision.

From that point, the students were fully responsible for generating the data with minimal assistance. They connected the devices to the computers, activated the relevant programs, loaded the priming mix (dubbed ‘fuel’) and the DNA libraries onto the flow cells, and launched the sequencing run using MinKnow. Once data was generated, they monitored the progress of the sequencing run. After checking quality measures, the sequencers were left unattended for 48 hours to generate data according to the ONT protocol.

After data generation, we instructed the students to complete an assignment, which was divided into two milestones (supplemental 5 and 6). The first milestone was reporting the technical performance of the MinION sequencer, such as the total reads, the read length distribution, and the read quality scores. The second milestone focused on an actual scientific problem the students tried to solve with the device (see next). For each milestone, the students had to submit a written report and a GitHub link to their code (an example: https://github.com/dspeyer/ubiq_genome). Each hackathon concluded with a 10-minute talk by each group.

Lessons learned from conducting hackathons:
- **Prepare spare parts:** During the hackathon, there is little time to troubleshoot. Over the 10 intended MinION runs (five groups over two hackathons), we experienced multiple technical difficulties. Three flowcells had an insufficient pore number (<51) and had to be replaced. In another event, a computer failed to connect with any MinION despite a working USB 3.0 port. It is therefore crucial to anticipate scenarios of failure and have spare parts (i.e. computers, flow cells, fuel mix, and DNA library).
- **Consider back-up data:** As part of testing our hackathon setting, we sequenced some of the DNA libraries with the MinION before the actual event. The data generated from these tests was kept to have a contingency plan in case none of the MinIONs worked at the time of the hackathon. This way students would still have data to analyze, and the course progression would not be jeopardized. While fortunately, we did not have to use this data in our cases, we encourage MinION hackathon organizers to consider this option.
- **Expect variability in the amount of data**: The yield of the MinION sequencers was variable between runs. The experimental design and the questions posed during each hackathon should be compatible with both a low and a high sequencing yield.
- **Locate appropriate computers:** One of our main challenges was to procure five computers that fell within ONT specifications. Our department is almost entirely Mac-based whereas the current ONT specification requires a Microsoft Windows computer. We tried installing Windows virtual machines on our Macintosh computers but found this solution unreliable presumably due to the fast data transmission rates of the sequencers. The students’ computers also fell short of the specifications required by ONT, such as having a solid-state drive. We propose MinION hackathon organizers to keep in mind that locating multiple appropriate computers can be a time-demanding task.
- **Network was not a problem:** ONT sequencing requires an Internet connection for base-calling. We connected the five computers to a regular network hub using a standard Ethernet protocol. We did not experience any issues.
- **Use free tools for data transfer:** MinION sequencing can result in large data folders. We looked for a free program to automatically transfer the data 48 hours after the start of the run from the sequencing laptop to the students’ computers. Cloud-based products, such as Dropbox, do not support synchronizing this amount of data with their free accounts. As an alternative, we used the free version of BitTorrent Sync, which was found to be apt for this purpose. This program allows sharing of files over the P2P BitTorrrent network without a size limit. Bit-Torrent can be pre-installed on the workstation and can be synchronized with the student’s personal computer by exchanging a folder-specific key. This solution for large files can be set up within a few minutes and prevents technical challenges.
- **Tune student expectations:** At the beginning of each hackathon, we reminded the students about the experimental nature of this event. We communicated clearly that they should anticipate technical issues and that we will be surprised if everything will go smoothly. This helped to reduce frustration for students, who are accustomed to interacting with mature technology in day-to-day life.

## Hackathon project 1

### Snack to sequence

The first Hackathon was called “From snack to sequence”. It was inspired by several food scandals such as the horsemeat containing ready-made meals that were labeled as beef throughout Europe in 2013 as well as the revelation that a number of NY sushi restaurants claimed to be selling a white tuna while in reality serving escolar. Based on this issue, we wanted to introduce students to the identification of species in different food items.

We prepared five sequencing libraries from dishes purchased at local NY restaurants as well as from raw food products that were purchased at a supermarket. The DNA libraries were a mix of multiple ingredients (like raw beef and tomato). We set out to address the following questions with the students: a) Can you identify the species in a food sample using MinION sequencing, without prior knowledge? b) Can you quantify the composition of the different ingredients? c) What is the minimal sequencing runtime required to detect the ingredients of the sample?

After generating the data in the hackathons, we devoted the next class to exploring a diverse number of sequencing algorithms that could be used for species identification. Importantly, Oxford Nanopore’s ‘What’s In My Pot’ species identification workflow does not support eukaryotic sample identification (Juul et al. 2015) and the students had to find alternatives. The consensus among the students of the class was that a BLAST search is the best option for identification.

Most groups were able to identify the species within the dish. One interesting discussion resulted from the two groups that sequenced samples putatively containing beef. The top BLAST hit was for bighorn sheep (*Ovis canadensis)*, whereas the domesticated sheep *(Ovis aries)* or cow *(Bos taurus)* was returned with lower alignment quality values. The identification of bighorn sheep was suspicious, since this animal is not domesticated. Cow is part of the *Bovidae* family, as are the bighorn and domesticated sheep. The students reasoned that the sample could be from a family member and selected the domesticated sheep as the most likely candidate. Another interesting finding was that the raw beef samples had DNA traces of infectious *Babesia bigemina* and cattle-vectored *Wuchereria bancrofti* and *Onchocerca ochengi* parasites with two or more reads per sample. These findings led to a vivid discussion on food safety.

Overall, this hackathon was academically apt for the level of the students. The only technical challenge the students repeatedly encountered was to BLAST a large number of query sequences using the program application-programming interface (API). They had to find creative solutions, such as mirroring the NIH BLAST to a private server and tweaking the input parameters to allow searching of a large number of long MinION reads.

### CSI Columbia

For the second hackathon, we explored the identification of individuals using ultra low coverage genome sequencing with the MinION. In forensics, DNA evidence identification relies on the analysis of the 13 well-characterized CODIS short tandem repeat (STR) loci (Kayser and de Knijff 2011). However, theoretical analysis has suggested that a small number (30–80) of common SNPs in linkage equilibrium are sufficient for positive identification (Lin, Owen, and Altman 2004). The aim of this Hackathon was to test whether it will be possible to identify a person using MinION shotgun sequencing with an extremely shallow coverage.

Two groups sequenced a DNA library prepared from genomic DNA from Craig Venter (Levy et al. 2007), one group sequenced a CEU HapMap sample, and two groups sequenced the genomic DNA of one of the authors (YE). Students were given very minimal information compared to the previous hackathon. They did not know whose genome they sequenced but were instructed to communicate with us to collect hints. We also encouraged the students to test additional methods to identify the person, such as examining the mitochondria haplogroup, the sex of the person, and estimating his or her ancestry.

Technically, the students generated data that covered ~5000 to ~29000 common SNPs. To increase the chance of uncovering the identity we hinted the student about the possible genomes that they sequenced to reduce their search space to about 1000 genomes.

The students found this assignment much more challenging than the previous one. Of the five groups, one was able to correctly identify the input sample (Craig Venter). The students tried an impressive set of tools but their main challenge was data wrangling. They had to convert their data to various formats in order to test different tools just to realize that the tools do not perform as expected or poorly documented, wasting a significant amount of time. Interestingly, some of the undergraduate students told us later that this was the first time they were exposed to an open-ended real-world research problem and that this task gave them a better understanding of academic research. The students also suggested that more discussions between the groups during the hackathon could have helped to solve some of the technical problems. This can be done using online communication tools (like Facebook or a Piazza website).

### Questionnaire

We also sought to more quantitatively learn about the views of students with respect to genomics and mobile sequencing. We asked them to answer a questionnaire before the first hackathon, when the students were exposed only to the theory of sequencing and its applications, and then three weeks later, after the completion of the first hackathon.

While our sample size is too small to draw statistical conclusions, we did learn from the trends in the answers. The hackathons seemed to have shaped a more realistic view of the technical challenges inherent to genomic applications. For instance, for the question “How long do you think it takes from sample preparation to sequencing results using MinION?”, about 70% of the students answered ‘one hour’ (or less) before the hackathon; after the hackathon, only 30% of the students thought it would take one hour. After the hackathons, students also thought that it would take more time for mobile sequencers to be used for health tracking by the general public and suggested lower monetary value for home sequencing applications. We did not observe changes before and after the hackathon for ethical issues such as “Do you think it is ethical to sequence hair found on the street?” or “do you think getting your genome sequenced is safe?” These trends suggest that the hackathon mainly shaped the students’ technical understanding and demonstrated the value of hands-on experience to facilitate realistic views of the challenges of new technologies.

## Concluding remarks

Mobile sequencing in the classroom proved to be a useful method for teaching students about the cross-disciplinary field of genomics. They enable an integrated view of the challenges in genomics. These devices are relatively inexpensive and do not require complicated equipment or designated lab space to be operated. As such, they dramatically reduce the barrier of classroom integration compared to other sequencing technologies.

The main focus of this manuscript was the integration of mobile sequencing as part of the higher education system (undergraduate and post-graduate). However, we also see the potential of integrating these devices in high school STEM curricula and enrichment programs. Such activities can expose pupils early in their training to the fascinating world of DNA and serve as an educational springboard to study other disciplines such as math, computer science, and chemistry. We hope that the resources and experience outlined in this manuscript will help to facilitate the advent of these programs.

## Acknowledgments

We thank James Brayer, Michael Micorescu, and Zoe McDougall from Oxford Nanopore Technologies for technical support and for providing the reagents to the class. We also thank Dina Zielinski for useful comments and discussions. Y.E. holds a Career Award at the Scientific Interface from the Burroughs Wellcome Fund. This work was partially supported by a National Institute of Justice (NIJ) grant 2014-DN-BX-K089 (Y.E. and S.Z.).

## Supplemental Note 1 Class website information

2015 Syllabus
| Week | Date | Topic |
| --- | --- | --- |
| 1 | 11-Sep | Intro to DNA and DNA sequencing technologies |
| 2 | 18-Sep | Applications of DNA sequencing: Human genetics |
| 3 | 25-Sep | Applications of DNA sequencing technologies: Metagenomics |
| 4 | 2-Oct | Application of DNA sequencing: Forensics |
| 5 | 9-Oct | Mobile health |
| 6 | 16-Oct | Mobile sequencing |
| 7 | 23-Oct | Hackaton I: "From Snack to Sequence" |
| 8 | 30-Oct | Analysis pipelines(1) |
| 9 | 6-Nov | Presentations for hackton I |
| 10 | 13-Nov | Hackaton II: "CSI Columbia" |
| 11 | 20-Nov | Analysis pipelines (2) |
| 12 | 27-Nov | [No class] |
| 13 | 4-Dec | Presentations for hackton II |
| 14 | 11-Dec | DNA as storage devices or computers |

### Assignments and grading

#### Reading assignments

You are expected to read the paper and understand the main concepts and terms before the class.

#### Presentations

The class has a few lessons that include team presentations. The length of each presentation is 10min and will be delivered by one member of the team. To encourage fairness and participation, the presenter will be randomly selected at the beginning of the presentation.

#### Coding/Written assignments

Teams are expected to code their own assignments. It is OK to brainstorm high-level ideas with other teams. It is OK to consult online forums. However, the submitted code should be fully written by members of the team. No exceptions. To maximize impact, all code should be submitted under the GNUv2 license.

### Grades

- Participation in class discussions: 25%
- Hackathon1: 25% (10% presentation + 15% code submission)
- Hackathon2: 25% (10% presentation + 15% code submission)
- Final project: 25%

## Supplemental Note 2

Protocol for setting up a Hackathon using the MinION:

### Sample preparation for MinIONs

For “Snack to Sequence”: Prior to the hackathon, food ingredients for which the genome is included in the NCBI database were selected from the supermarket. The DNA of seven food products was isolated using standard procedures, including four raw products (tomato, kale, beef, bacon) and three that were cooked (salmon, shrimp, and mussel). The DNA samples were fragmented using Covaris g-TUBEs and further processed by end-repair and dA tailing using the NEBNext Ultra II End Repair/dA-tailing Module. The sequencing libraries were prepared according to version 6 of Oxford Nanopore Technologies Protocol (MAP006). DNA concentration was measured by Qubit Fluorometric Quantitation (Thermo Fisher). The composition of the DNA library samples prepared for the 5 groups is indicated below:

| Students | Ingredient | ratio |
| --- | --- | --- |
| group1 | beef | 80% |
|  | tomato | 20% |
| group2 | bacon | 80% |
|  | salmon | 20% |
| group3 | kale | 10% |
|  | mussel | 60% |
|  | shrimp | 30% |
| group4 | beef | 20% |
|  | tomato | 80% |
| group5 | bacon | 20% |
|  | salmon | 80% |

For “CSI Columbia”, DNA samples for Craig Venter (NS12911) and the HapMap consortium (NA12890) were purchased from the Coriell Institute. Genomic DNA from Yaniv Erlich was extracted from saliva using Quiagen QIAamp^®^ DNA Blood Mini Kit. DNA samples were enriched for fragment size by gel electrophoresis and isolation of fragments >3kb (Quaigen^®^ Gel extraction kit). These fragments were sheared using Covaris g-TUBE size ~6 kb. The generation of the DNA libraries was done as described for “Snack to Sequence”.

Two days prior to the hackathon the library was prepared and tested for functionality. The libraries can be stored in 4^°^C up to 3 days.

### MinIONs in the classroom

In addition to the MkI MinIONs, ONT sponsored the class by supplying fresh flow cells a few days before the start of the hackathon. The MinION flow cells contain a port where the sample is applied. This port is the opening to a channel that pushes the applied reagents towards the pore-chamber. A new flow cell contains a small quantity of air at the opening that needs to be removed otherwise it will be pushed toward the pore-chamber and perturb the accessibility of the DNA strands to the pores. Removal of the air bubble requires attention and possibly help from instructors.

On the hackathon day we pre-prepared the fuel mix (required for priming the MinION flow cell) one hour prior the hackathon start. The library mix was freshly prepared 10 minutes before loading of the samples.

### Computers and software

#### MinKnow

Oxford Nanopore Technologies provided the class with 5 MkI MinIONs. The software interface MinKnow runs specifically on Windows computer systems at the time of writing. The Windows computer must have a USB 3.0 port and sufficient storage space for the data generated (we used up to 40 Gbytes). The computer requires installation of MinKnow and USB 3.0 driver. Running the MinION configuration menu in MinKnow tests correct installation of the software. MinKnow ran using the menu “MAP_48Hr_Sequencing_Run_SQK_MAP006”.

Computers should be able to run for up to 48 hours. We advise verification of the computer sleep-mode before starting a run.

#### Basecalling

Basecalling is performed via the cloud service Metrichor. Metrichor can be downloaded and installed on the workstation. Installation requires presetting the directory used for uploading raw squiggle data and downloading the base-called sequencing data. Moreover, the proper setting of the user key allows for following the sequencing run yield in real time. For both hackathons the “2D basecalling for SQK-MAP006” recipe was used, which requires the base-called read to be 2D (i.e. successful passing of both template and complement) with an average Quality Score >9.

#### Data Transfer

The base-called data was synchronized to the student computers via BitTorrent. Selecting a folder creates a folder-specific key, which can be provided to the students. Via a manual connection on the personal laptop, the student can paste the key. This instantly starts synchronization between workstation and personal laptop. Before the hackathon it is advisable to notify students that they require a minimum of ~20 GB of storage on their computers (this number depends on the productivity of the flow cell, the DNA library and the duration of the run). A possible alternative is the use of an external hard-drive. During both hackathons, one computer would not connect to the MinION and one computer stopped generating data after going into sleep mode. These issues should be considered when setting up the operating system.

## Supplemental Note 3

### Assignment | Snack to Sequence

Please **read** and **follow** these instructions:

**October 25^th^ 11:59pm:** to notify that you were able to synchronize you computer using BitTorrent.

**October 30^th^ Noon:** first submission ‘Quality Assessment MinION reads’.

**November 6^th^ Noon:** second submission ‘Snack to sequence pipeline’.

**November 6^th^**: presentations ‘Snack to sequence’. Each team will have 10 min to present. Two presenting members will be selected randomly at the beginning of class.

**For both submissions submit to** with subject line [Hackathon#1, assignment number, groupID]

- Submit the written portion of your homework.
- Submit all code written by the group in their original extensions. Any programming language is acceptable, although Python is preferred.
- Your code should have a <u>GPLv2</u> license.
- Submission can be done by emailing a link to a git repo. If the team does not know how to use git, they can email a tar ball of the code.

#### Late submission policy

Failures to meet deadlines will result in 25% grade reduction for each late day.

#### Assignment 1 Quality Assessment MinION reads

Convert the data from fast5 files to fasta files and fastq using ‘Poretools’.

Analyze the following parameters:

1. Calculate the number of 1D and 2D reads classified as ‘failed’ versus the number of 1D and 2D reads classified as ‘passed’. Calculate the fraction of reads that are 2D called in both the ‘pass’ and ‘fail’ folders.
2. Calculate average reads per active channel (remember you wrote the number of active pores down for group 1 during the hackathon). Which channel in the flowcell produced the most reads? How many?
3. Plot the cumulative <u>nucleotides</u> sequenced as a function of time for both ‘failed’ and ‘passed’ 2D reads in separate graphs.
4. How many hours would you have to sequence in order to cover the human genome once? (Only using 2D reads that passed the quality filters.)
5. Metrichor uses a base-calling algorithm that gives the accuracy with which the sequencing platform could identify the particular base (FASTQ files include quality scores). The quality scores are based on ASCII, which is a character encoding system that maps a number to a character. Calculate the base-calling quality mean and standard deviation for 2D reads for both the ‘failed’ and ‘passed’ reads, and compare using a student’s t-test. In addition, compare the median base quality for the ‘passed’ 2D reads from the first hour with the median base quality of the last hour of that same sequencing run. Briefly comment on your results.
6. Plot a histogram the length distribution of 1D reads (template and complement) and the 2D reads in the failed folder. Do the same for the ‘passed’ reads.
7. Identify the longest read you obtained for: template, complement, and 2D from the passed reads. State the number of nucleotides for each.
8. Analyze whether there is a correlation between sequence length and timing of a DNA strand passing through the pore. Plot the obtained sequence length over time for 2D reads. Briefly comment on your results.
9. Plot the pace of the strand sequencing (sequence length per duration in pore) for 2D reads classified as ‘failed’ versus reads classified as ‘passed’.
10. Define the nucleotide composition of both 2D sequences classified as ‘passed’ and as ‘failed’ (calculate the percentage of G,C,T, and As in the results).
11. Build a model that predicts the time to sequence a segment based on the input sequence. You can use any type of classifier or features you want. Report the cross correlation r^^^2.

Your submitted code should be able to replicate the output of your report. Document your code. You can write a separate program for each question. The naming of your code should be groupX_report1_questionY, where X is your group number and Y is the question number

#### Assignment 2 Snack to sequence pipeline

1. **Develop an analysis pipeline for:**

- Identification of food ingredients in your sample.
- Once you identified the ingredients, quantify the ratios in your dish.
- Do you find any bacteria? *Bonus points* if you can develop a simple web interface/app that takes MinION reads and generates a visual real time analysis.
2. After how many minutes in a MinION run would you be able to state what the composition of your food was and in what ratio?
3. Filter the sequences for one of the food components.

- From those sequences, make a confusion matrix. A confusion matrix takes a known reference sequence, and tests the classification of your reads.
- Based on the alignments obtained, filter only the deletions and insertions. Of the deletions and insertions found, calculate the size distribution and the nucleotide composition.

**Presentation:** Snack to sequence pipeline

The presentation should include the following items:

1. Report the output of the sequencer and the longest read.
2. Present the number of errors and quality of the sample.
3. Present the classifier and features of reading speed and performance.
4. Strategy for identifying Sophie’s food.
5. What biological ingredients you find in her food.
6. Suggestion for a follow up question

## Supplemental Note 4

### Hackathon #2 | CSI Columbia

#### Assignment 3: QC and alignment

1. Calculate the number of 1D and 2D reads classified as ‘failed’ versus the number of 1D and 2D reads classified as ‘passed’. Calculate the fraction of reads that are 2D called in both the ‘passed’ and ‘failed’ folders.
2. Calculate average reads per active channel (remember you wrote the number of active pores down for group 1 during the hackathon). Which channel in the flow-cell produced the most reads? How many?
3. Plot the cumulative <u>nucleotides</u> sequenced as a function of time for both ‘failed’ and ‘passed’ 2D reads in separate graphs.
4. Plot a histogram of the length distribution of 1D reads (template and complement) and the 2D reads in the failed folder. Do the same for the ‘passed’ reads.
5. 2D from the passed and failed reads. State the number of nucleotides for each.
6. Align the reads to the human genome (hg19) using BWA-MEM ONT. Report how many reads aligned.
7. Report a confusion matrix of your reads
8. Open question: suggest three strategies to reduce the number of errors in the reads.

Your submitted code should be able to replicate the output for your report. Document your code. You can write a separate program for each question. The naming of your code should be groupX_report1_questionY, where X is your group number and Y is the number of question (or questions).

#### Assignment 4

Try to find as much as you can about the person you sequenced:

- Who is this person?
- Ancestry?
- Sex?
- Phenotypic traits?
- Go wild!

Helpful links:

- https://genome.ucsc.edu/
- http://www.1000genomes.org/
- https://imputationserver.sph.umich.edu/index.html
- http://pritchardlab.stanford.edu/structure.html
- https://opensnp.org/

#### Presentation: CSI Columbia

The presentation should include the following items:

1. Report the output of the sequencer and the longest read.
2. Present QC of the data.
3. Present the error frequency, and confusion matrix.
4. Present the pipeline used to identify the traits.
5. Present all traits you could find.

